# Feedback-related potentials and oscillations during trial and error learning in Parkinson’s disease

**DOI:** 10.1101/2021.04.05.438433

**Authors:** Laura Viñales, Emmanuel Procyk, René Quilodran

**Author notes:** Author Contributions*: LV, EP RQ Design Research; LV and RQ Acquired Data, LV Analyzed data, LV and EP Wrote the Paper.

## Abstract

Electrophysiological markers of performance monitoring are thought to reflect functioning of dedicated neural networks and neuromodulatory systems. Whether and how these markers are altered in neurological diseases and whether they can reflect particular cognitive deficits remains to be confirmed. Here we first tested whether the frontal medial feedback-related potential, evoked during a trial and error learning task, is changed in early Parkinson’s disease patients compared to control subjects. The potential was not changed in amplitude and discriminated negative and positive feedback as in controls. Feedback-related markers in Parkinson’s patients also appeared in time-frequency analyses, unaltered in theta (3-7 Hz) band but reduced in beta (20-30 Hz) oscillations for positive feedback. Beta oscillations power appeared to be dramatically globally reduced during the task. Overall, our results show that Beta oscillation markers of performance monitoring captured by EEG are selectively altered in Parkinson’s disease patients, and that they are accompanied by changes in task-related oscillatory dynamics.

**Significance Statement:** Frontal neural activity evoked by outcomes reveal the functioning of neural systems devoted to flexible behaviours. Modulations of such activity in Parkinson’s disease (PD) patients could reflect specific alterations of neural systems and cognitive processing. The goal is to evaluate whether such activity can serve as markers of the disease. Here, using EEG and a trial and error learning protocol, we show that mid-frontal midline performance feedback-related potentials for different types of feedback were similar in controls and early diagnosed PD patients. However, task-related oscillations revealed alterations in the beta range accompanied by more global beta activity alteration in PD compared to controls subjects. This study provides data relevant to the search for non-motor biomarkers in early stages of PD.

## Introduction

Besides the cardinal clinical features of Parkinson’s disease (PD), including dysfunctions of the somatomotor system, non-motor symptoms develop in the early stages of PD and might often precede motor disabilities (Simuni and Sethi, 2008; Rodriguez-Oroz et al., 2009; Del Tredici and Braak, 2012). These symptoms encompass impairments of executive functions, and neuropsychiatric (e.g. apathy, depression) or sleep disorders. Executive impairments, are important predictors of quality of life (Schrag et al., 2000), and are present in a large proportion of patients with PD (20-40%) at initial diagnosis although they are often overshadowed by motor symptoms (Foltynie et al., 2004; Muslimovic et al., 2005). Disabilities for planning and adapting behavior, or evaluating outcomes of actions are some of cognitive dysfunctions observed in PD (Owen et al., 1992; Cools et al., 2001; Dirnberger et al., 2005). Identification of markers of Parkinson’s disease would be beneficial for early detection and early therapeutic interventions, and their potential relationships to behavioral or cognitive alteration would help understand the evolution of the disease.

Two well studied markers of performance monitoring are the error-related negativity (ERN), an event-related potential evoked at the execution of incorrect responses in reaction time tasks, and the feedback-related negativity (FRN) observed after the presentation of a sensory feedback signaling performance (Sallet et al., 2013; Ullsperger et al., 2014). One possible central source of the FRN is the midcingulate cortex (MCC), a cortical region receiving important mesocortical dopaminergic input and involved in feedback processing (Williams, 1998; Procyk et al., 2016). The Reinforcement Learning theory of the ERN causally links these markers to the dopaminergic system and the MCC (Holroyd and Coles, 2002). Recordings in monkeys MCC revealed that some aspects of feedback-related single unit activity are modulated in accordance with reinforcement-learning like rules, but others seem to relate more to feedback categorizations and the expression of switch-stay behavioral strategies (Matsumoto et al., 2007; Quilodran et al., 2008). One assumption is that performance monitoring signals (FRN or ERN) correlate with the state of dopaminergic transmission (Jocham and Ullsperger, 2009). This has been tested pharmacologically in non-human primates for FRN (Vezoli and Procyk, 2009; Wilson et al., 2016) and in humans (Zirnheld et al., 2004). In a progressive and pre-symptomatic monkey model of Parkinson’s disease, the FRN was the most altered marker of the lesion, although this effect was found in the absence of behavioral deficit in a deterministic trial and error learning task (Wilson et al., 2016). Several EEG studies have demonstrated amplitude attenuation of the ERN in PD patients (Falkenstein et al., 2001; Ito and Kitagawa, 2006; Willemssen et al., 2009a) whereas other studies did not, depending on the tested group or medication status (Holroyd et al., 2002; Stemmer et al., 2007). Note importantly, that the FRN can be associated to both negative and positive feedback of performance (Sallet et al., 2013). However, to our knowledge there are only rare studies investigating FRN modulations in PD patients. One recent study showed that a fronto-central positivity to positive Feedback was diminished in PD patients compared to controls in a probabilistic two alternative forced choice task (Brown et al., 2020). The reward-related potential did not vary with ON-OFF status of medications, but was more reduced for early compared to later diagnosed patients. Finally, the FRN for negative outcomes (no reward) did not vary between subject groups.

Other relevant neurophysiological markers are oscillation power evoked or induced during tasks and that might reflect basic mechanisms of communication, processing, and might contribute to plasticity (Buzsáki and Draguhn, 2004). Theta and beta oscillations have been observed after negative and/or positive outcomes or feedback (Cohen et al., 2007; Marco-Pallares et al., 2008; Marco-Pallarés et al., 2009; De Pascalis et al., 2010b; Doñamayor et al., 2011; van de Vijver et al., 2011; HajiHosseini et al., 2012; HajiHosseini and Holroyd, 2015). Investigations have shown important alteration of subcortical beta oscillations in PD patients in relation to sensorimotor events but also to reinforcements (Schroll et al., 2018). In addition, alterations of oscillation power measured by EEG in PD patients have been observed at low frequency (delta and theta bands) during cognitive tasks and compared to age-matched healthy controls, suggesting that altered signal in those bands (especially on the frontal midline) might be used as a biomarker of dysfunction in Parkinson’s disease (Schmiedt-Fehr et al., 2007; Singh et al., 2018).

The relationships between FRN, outcome-related oscillations, positive and negative feedback processing and Parkinson’s disease thus remain unclear. Because of the asymmetric nature of dopaminergic single unit firing for negative (inhibition) and positive (activation) prediction errors (Schultz, 2007), it was postulated that the effect of positive prediction errors is stronger on FRN than negative prediction errors (Holroyd et al., 2008). However, studies of single unit activity in MCC have repeatedly shown that single unit populations contributing to negative versus positive outcome processing are separated (Matsumoto et al., 2007; Quilodran et al., 2008; Kennerley et al., 2009). Alternatively, the FRN (and its variance explained by outcome value) is also conceived as a reward positivity acting on top of, and reducing, a generic negative deflection observed for other events including negative feedback (Holroyd et al., 2008; Proudfit, 2015). This alternative would indeed suggest two separate sources.

The alteration of these features in Parkinson’s disease is poorly described. Here we used a behavioral protocol involving robust adaptive components (trial and error learning task) to test the potential modifications of FRN and oscillatory activity in the beta and theta bands by altered dopaminergic states in Parkinson’s disease. We took advantage of some properties of the protocol (various forms of negative and positive feedback) to study the modulations of FRN and oscillations in Parkinson’s disease patients compared to control subjects, and to test whether the altered neural states in patients would differentially influence frontal responses to positive and negative feedback.

## Methodology

### Participants

Seventeen patients with idiopathic PD and fifteen age-matched healthy controls participated in the study. All participants gave signed informed consent according to the declaration of Helsinki after they were introduced to the purpose of the study and after the protocol was explained to them. The study was performed with the approval by the review boards and the School of Medicine’s ethics committees.

The patients were recruited at the Department of Neurology and Parkinson’s disease Health Center, and the diagnosis of PD was based on the clinical criteria of the United Kingdom Parkinson Disease Society Brain Bank (UPKD) (Daniel, 1993). The mean age of the patients was 64.9 +/-7.1 years (+/-SD). Diagnosis of PD was made on average 4.9 years before EEG recordings. All patients were receiving L-dopa therapy and were selected for being at clinical stage 1-3 using Hoehn and Yahr classification (HYS) i.e. they were at the initial stages of the disease (Hoehn and Yahr, 1967), mean 2.4 +/-0.6. Motor status and quality of life were determined by the motor subscale of the UPDRS (Fahn and Elthon, 1987; Goetz et al., 2008), with evaluation of 37.3+/-16.3, and PDQ-39 test (30+/-17.6). During experimental sessions, all patients were in OFF medication (medication was stopped on the evening prior to the experiment; the experiment was performed in the morning; medication was re-initiated after the test). Each patient’s medication was checked and doses of antiparkinsonian medication were converted to levodopa equivalent daily dose (LED) according to the algorithm used in Tomlinson and colleagues (Tomlinson et al., 2010).

None of the patients had dementia, history of cerebral infarction, pronounced tremor or presence of atherosclerosis. The mean age of control subjects was 65.9 +/-6.6 years (+/-SD). None of the selected control subjects had any history of neurological or psychiatric disorder (no history of dementia, no cognitive deficit tested ensuring ACE-R < 74/100, a BDI-II depression score < 18, and an apathy score (AS) < 14) and none of the subjects was taking drug affecting the central nervous system. All participants were right-handed. All subjects were assessed with a series of neuropsychological tests in a separate session before EEG recording to investigate participant’s cognitive and emotional status. Cognitive functions were estimated using the revised version of the Addenbrooke’s Cognitive Examination (ACE-R) (Larner and Mitchell, 2014), controls: 91.7 +/-5.5 and PD patients: 86.55 +/-6.6 (1-way ANOVA on Subject Group effect, F(1,30)=6.68, p = 0.015). Apathy and depression were evaluated using the Apathy scale (AS) (Starkstein et al., 1992), control: 4.2 +/-2.9 and PD patients: 11.1 +/-7.3 (1-way ANOVA, F(1,30)=13.06, p = 0.001), and the Beck Depression Inventory – Second Edition (BDI-II) (Beck et al., 1987), control: 10.5 +/-6.1 and PD patients 13.8 +/-9.1 (1-way ANOVA, F(1,30)=1.46, p = 0.26). The two groups of subjects thus differed in terms of apathy score, and cognitive and education level scores.

### Behavioral protocol

To assess executive functions and markers of behavioral adaptation we used electroencephalography (EEG) while participants were performing a modified version of a trial and error task. Each subject was comfortably seated in an armchair at 100 cm in front of a 21-inch monitor screen, on which visual targets were presented using E-Prime 2.0 Professional Edition (Psychology Software Tools, Pittsburg, PA, USA).

During the execution of the task, each subject had to search by trial and error which of four visual targets (filled disks, upper left, upper right, lower right, and lower left; **Fig. 1A**) simultaneously presented on a video screen was associated with a positive feedback. A trial started with the onset of a white central fixation point 1000ms before the onset of the four grey targets. Subjects were instructed to fixate the fixation point during the entire trial. 1300ms after the target onset presentation, all four targets turned white as a “Go Signal”. The subject could then respond by pressing one of four buttons on a response box and within a time limit of 2500ms. Subjects were asked to react as fast as possible to the “Go Signal”. At button press, all targets switched off and a delay of 1000ms elapsed before the presentation of a visual feedback lasting 700ms. Feedback corresponded to a central red or green square for negative and positive outcome, respectively. After each incorrect choice (INC) the subject could continue searching for the correct target. The discovery of the correct target was indicated by the presentation of the first green feedback (CO1), representing the end of the search period and the beginning of the repetition period. During the repetition period, subjects could repeat the correct choice for two trials (COR). A repetition ended with the onset of a central yellow circle, named “Signal-to-Change” (SC), presented 700ms after the feedback offset and for 1000ms. The SC stimulus indicated the end of the current problem, and thus the initiation of a new search for the following problem (see figure 1a). Each block was composed of 28 problems, and all participants made 2 blocks (performed in 2 separate sessions).

**Figure 1.**
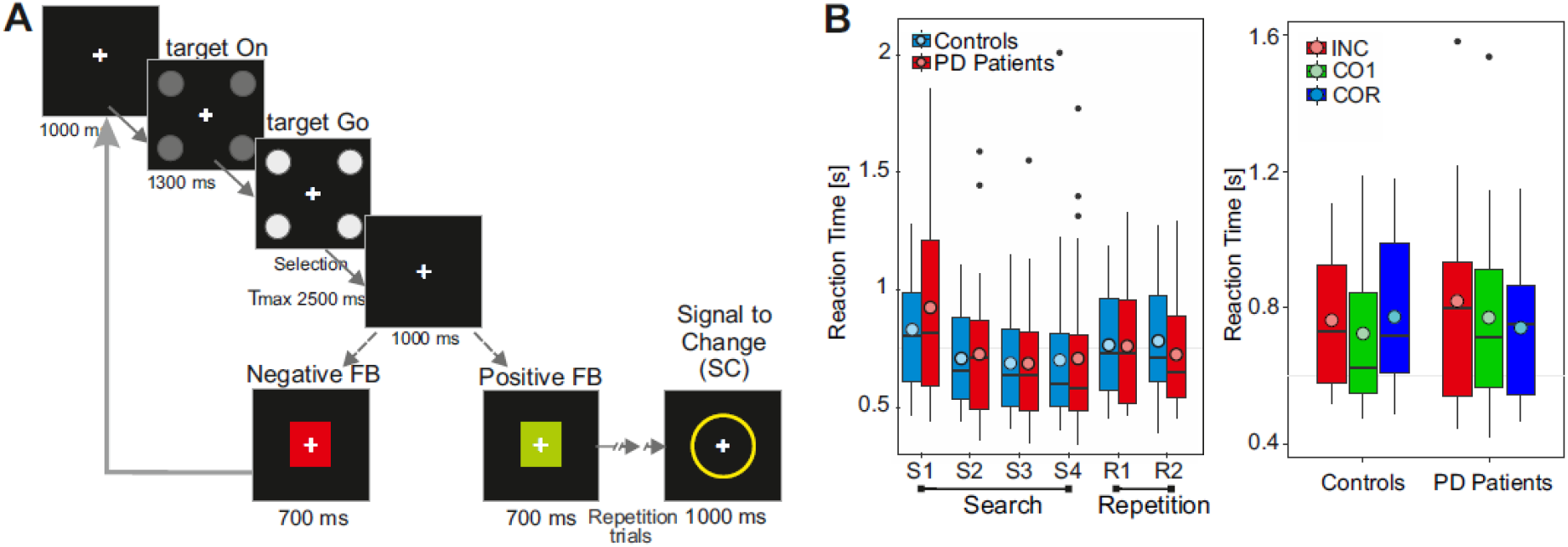
A. Behavioural task. Each trial starts with a fixation point, then the targets appear, showing the options to choose from. Target presentation is a signal to prepare the response. Then a GO signal is presented (“Target Go”, targets turn white), and the subject can select a target and touch the corresponding button in less than 2.5 sec. Feedback is delivered for 700ms - positive (green box) or negative (red box). The search period ends when the correct choice is made (1^st^ correct feedback). The repetition then starts, and the subject must select the same option in two more trials before the problem is completed and the signal to change appears indicating the start of a new search period. **B. Left**. Reaction Times PD Patients vs Control. Mean Reaction time box plot of PD patients (red) and controls (blue), the x-axis presents the choice instance, both in search and repetition. The circle represents the average reaction time per trial and subject group. **Right**. Reaction Times in Control and PD patients by feedback type. Mean Reaction time box plot for Incorrect Feedback (INC, red), first correct feedback (CO1, green) and correct feedback in repetition (COR, blue). The colored disks over boxes represent the mean values.

### Behavioral analysis

Performance in the trial and error task was quantified using : the number of unsolved problems, defined as problems where the subject did not complete the search or repetition periods; the number of *Repetition Errors*: correspond to errors committed only in the repetition period; the number of *Search errors* which are divided into: *memory errors*, defined as the selection in the search period of one target that was already selected, but not consecutively, and associated with a negative feedback (for example, the four targets being designated as 1 to 4, a choice sequence with a memory error would be: 3 2 3 1 …), and *Perseverance errors*, where despite receiving a negative feedback the same option is selected again (e.g. 3 2 2 1) .

Response times (i.e. time between the Go signal and the touch of a button) were analyzed and grouped in trials for the first, second, third and fourth search trials, and for the first and second repetition trials of each problem.

### EEG recordings and analyses

Each experimental recording was preceded by a practice to ensure that subjects (controls and patients) had understood the behavioral task. During task performance the electroencephalogram (EEG) was recorded using a 40 channels NuAmp Express System from 31 active Ag/AgCl electrodes using the standard extended 10-20 system: Fp1, Fp2, F7, F5, F3, Fz, F4, F6, F8, FC3, FCz, FC4, FT8, T3, C3, Cz, C4, T4, TP7, CP3, CPz, CP4, TP8, T5, P3, Pz, P4, T6, O1, Oz and O2. The horizontal EOG was recorded from 2 electrodes at the outer canthi of the eyes, and the vertical EOG was recorded from 2 electrodes above and below the left eye. The 40 monopolar channels NuAmps amplifier was used (Neuroscan Inc.). The forehead was used as ground. EEG and EOG data were sampled at 1000 Hz (Acquire, Neuroscan Inc.) with 0.1 to 100 Hz band-pass filter, and stored continuously on a PC hard-disk together with stimulus and response markers. Individual Ag/AgCl electrodes were adjusted until the impedance stayed below 5 KΩ.

### EEG Pre-processing

Continuous signal analyses were done using Matlab 2014 software (Matlab, The MathWorks Inc. Natick, MA, USA) and the Fieldtrip toolbox (Oostenveld et al., 2011). All EEG data were segmented in epochs from -1000 to 2000ms centered on the onset of feedback stimulus, with a large window to accommodate edge artifacts induced by wavelet convolution. We removed trials with signal amplitudes superior to 200µV. We applied a Butterworth filter with zero phase, between 1 and 60 Hz and a specific filter DFT to remove 50Hz. Eye-blink artifacts were corrected by using SOBI (Second Order Blind Identification). To represent the results, data were segmented between -100 and 600ms centered on feedback onset.

### FRN

A baseline correction was applied to all segments, by subtracting the average of the -300 to -100ms window for each trial. The FRN-amplitude was defined as the peak to peak amplitude difference between the negative peak detected in the 250 - 400ms time window and the preceding positive peak detected in the 150 - 300ms window at electrode Fz. Trials were then classified according to feedback type (INC, CO1, COR) and group (control, PD patients) to obtain the average signal.

The number of segments used for EEG analyses for the two groups were, for Parkinson disease patients group: INC, mean=79.6, std=11, min=56, max=96; CO1, mean=54.9 std=3.1, min=45, max=56; COR: mean=110.3 std=5.4, min=91, max=112, and for Control group: INC, mean=84.6 std=10.8, min=64, max=104; CO1, mean=55.3 std=2.3, min=47, max=56; COR, mean=108.9 std=6.6, min=88, max=112.

### Time frequency (TF) analyses

Each epoch was convolved with a set of Morlet wavelets, with the following parameters: number of cycles 6, width of Gaussian kernel 3. The frequency band of interest was 3 to 60Hz in 100 logarithmically spaced steps. A baseline correction was applied using the frequency average power from -300 to -100ms. The peak amplitudes of activity of the average resultant matrix for each subject was extracted in 2 windows of interest, theta band: 3 to 7Hz, in the windows 200 to 600ms post-feedback, and beta band: 15 to 30 Hz in the 300 to 600ms window post-feedback. The ROIs for time-frequency analyses were chosen based on our own experience of TF analyses with human and animal data as well as based on previously used ROIs. Frequency limits are usually around these limits: e.g. 1-8Hz in Lara and Wallis 2014, 3-9 Hz in Babapoor-Farrokhran et al. 2017, 4-8 Hz in Singh et al. 2018 and Narayanan et al 2013 for Theta (Narayanan et al., 2013; Lara and Wallis, 2014; Babapoor-Farrokhran et al., 2017; Singh et al., 2018). Regarding Beta band limits we chose the 15-30Hz limit to cover low and high beta ranges as several types of beta frequencies have been observed (e.g. see Stoll et al., 2016). We indeed separated in 2 different Beta bands, High (12-20Hz) and low (20-30Hz) in the continuous signal analyses presented in figure 4 (see below).

The time extents of ROIs were chosen according to frequency bands and based on visual inspection of time-frequency diagrams, covering the ensemble of phenomena observed post-feedback. We chose a longer window (200 to 600 ms) for the Theta band and a shorter one for Beta band (300 to 600 ms).

In addition, we performed power spectrum density analyses on the continuous recorded EEG for controls and PD patients. We used the same initial processing methodology as for time-frequency analyses, but taking the first 13 minutes in the task per subject as a continuous signal (13 minutes corresponding to the maximum duration of stable continuous signal, across all subjects), on which we applied a filter between 1 to 60Hz to then obtain the grand average per group, controls and PD patients. Finally cross-frequency correlation matrices were calculated from power spectral density for controls and PD patients (Llinás et al., 1999).

### Statistical analyses

All statistical analyses were performed using R (R Development Core Team, 2008). For behavioral analyses, each variable (number of unsolved problems, number of repetition error, number of memory error and number of perseverative error) was fitted using mixed generalized linear models with Poisson distribution with 2 (group: Patient, Control) x 2 (Block number: Block 1, Block 2) design. For response time analyses we performed 6 (Stage: 1^st^ Search, 2^nd^ Search, 3th Search, 4^th^ Search, 1^st^ Repetition, 2^nd^ Repetition) x 2 (group: Patient, Control) x 2 (Block number: Block 1, Block 2) repeated-measures ANOVAs. The reaction time measured in trials with different feedback types were analyzed using a 3 (Feedback: INC, CO1, COR) x 2 (group: Patient, Control) repeated-measures ANOVA. For the FRN amplitude analyses we performed 3 (Feedback: INC, CO1, COR) x 2 (group: Patient, Control) repeated-measures ANOVA. The effect of condition (Control vs Patient) was directly evaluated with 1-way ANOVAs.

In time-frequency analyses, measures were tested against baseline values through 1-way ANOVAs for PD patients and controls for the 3 feedback types. The direct contrast between measures in Controls and Parkinson groups was also tested independently for each feedback. In both cases multiple measures were accounted for by using FDR correction.

Statistical analyses were carried out with a significance threshold of p = 0.05. Sphericity was tested prior to running repeated measures ANOVA using a Mauchly’s test of sphericity. The Greenhouse-Geisser epsilon was used to correct for possible violation of the sphericity assumption. We report p-value after the correction.

## Results

### Behavior

Results were obtained after the analysis of 15 control subjects in 30 recording sessions and 17 PD patients in 34 recording sessions. PD patients performed significantly more unresolved problems (mixed Poisson glm, group effect: Deviance(1,62) = 54.7, p = 0.0055) and repetition errors (group effect: Dev (1,62) = 131.9, p = 4.11e-5) than controls, but the number of memory errors were not significantly different between subject groups, and did not vary between blocks. The frequency of perseverance error differed between group and increased in the second block of the session (Poisson glm, Group x Block interaction, Dev (1,60) =128.5, p = 0.024).

To study RTs of PD patients and control subjects we focused on search and repetition trials from correctly solved problems. We sorted trials according to their rank in the problem (First to fourth trials in the search periods and first and second in the repetition periods). This provides a view of the average dynamic of RTs during a problem (**Fig. 1B**). RTs were not different between groups (PD patients and control subjects) or between blocks (1^st^ versus 2^nd^ block). For both groups, RTs were longer in the first search trial and decreased over the search period (mixed glm, group x block x trail-rank, F(5,330)=14.07, p < .0001).

### Feedback-related negativity

The feedback-related potential is modulated by feedback valence and outcome expectancies (Cohen et al., 2007; Sallet et al., 2013). Because its modulation might depend on the dopaminergic system, we looked for changes between controls and PD patients. The electrophysiological responses for the different types of feedback (negative feedback: INC, first positive feedback: CO1 and positive feedback in repetitions: COR) for control and PD patients at electrode Fz are shown in **Fig. 2A**. The distributions of individual FRN peak amplitude (negative – positive peak, see methods) per group are shown in **Fig. 2B**. We found no interaction between feedback type (INC, CO1, COR) and group (control, PD patient) (type x group, F(2,60) = 0.62, p = 0.54), but a main effect of feedback type (type, F(2,60) = 34.7, p < .0001) on the FRN (**Fig. 2B**). On the Scalp topography, for both groups and the three types of feedback, the negative frontal negativity is clearly observed in control subjects for INC and CO1 **(Fig. 2C**). An analysis across midline electrodes showed an effect of electrode on the difference of amplitude between INC and COR and between CO1 and COR, but no effect of group (ANOVA, electrode x group, CO1-COR: electrode F=4.86, p<0.01, group F=3.4, p=0.06, interaction: ns; INC-COR: electrode F=5.53, P<0.001, group F=2.58, p0.1, interaction: ns). In controls, the FRN observed in response to feedback at Fz in INC and CO1 trials was larger than in COR trials. PD patients only the FRN at CO1 mean amplitude differed from COR (Tukey Post-hoc statistics, Bonferroni adjusted, Control_COR vs Control_CO1: Estimate= -2.94, Std Error=0.52, z-value= -5.66, p-value=2.3e-7; Control_COR vs Control_INC: Estimate=-1.93, Std Error=0.52, z-value=3.71, p-value=0.003 ; PD_COR vs PD_CO1: Estimate=-2.99, Std Error=0.49, z-value=-6.12, p-value=1.44e-8). However, directly contrasting FRN measures for the different feedback types in Controls vs. Patients revealed that FRN measures were not impacted by condition: INC F(1,30) = 1.31, p=0.26; CO1: F(1,30) = 0.1, p = 0.76 and COR: F(1,30) = 0.25, p = 0.62. Thus, the amplitude and pattern of FRNs was present in the PD group of subjects like in Controls.

**Figure 2.**
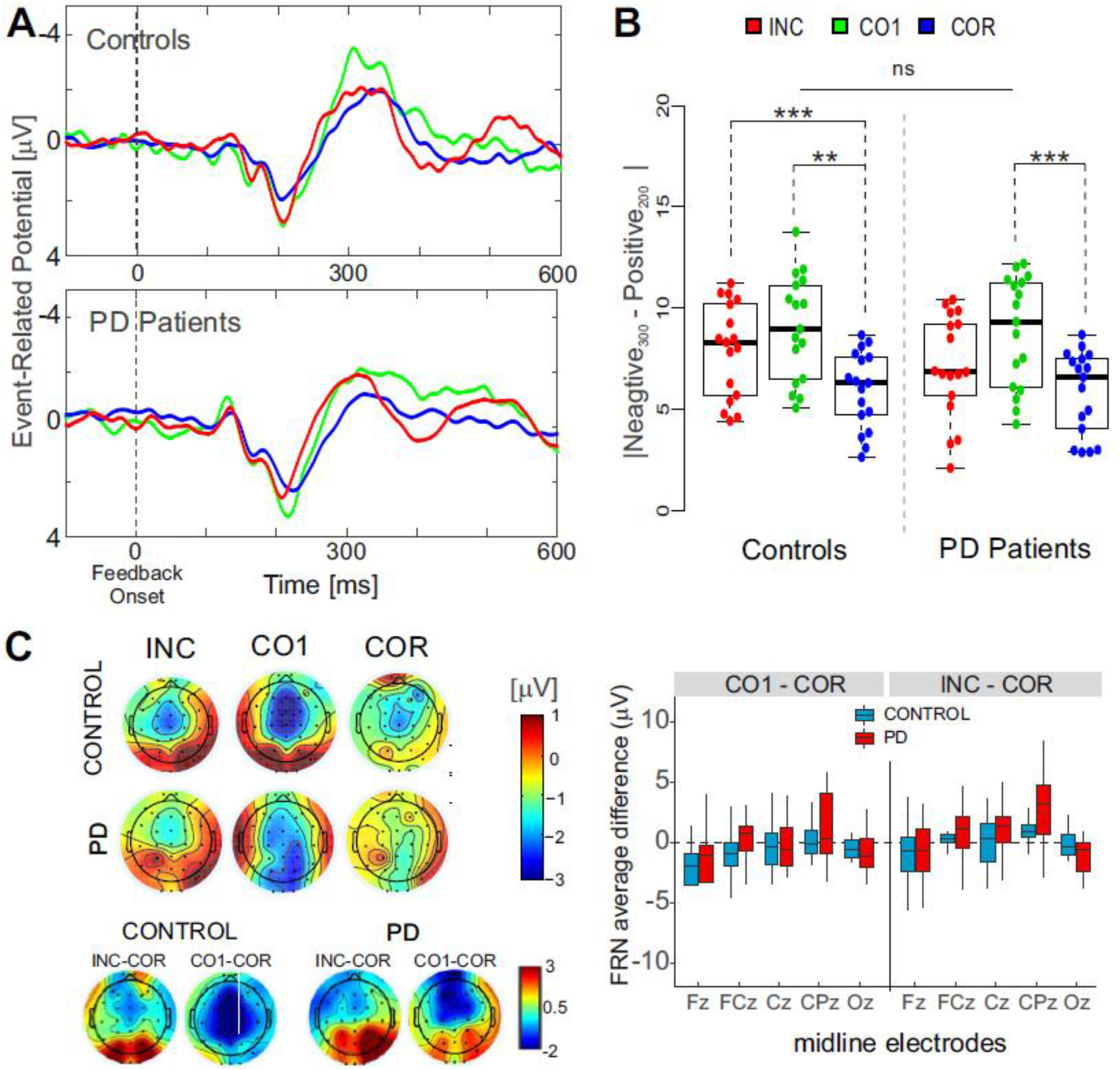
A. Grand-average electrophysiological feedback responses for INC, CO1 and COR trials in controls subjects and PD patients at electrode Fz. ERP are shown for incorrect response (INC, in red), first correct response (CO1, in green), correct response in repetition (COR, in blue), and are aligned on feedback onset. **B**. FRN amplitude. Distribution of differences between the negative peak and preceding positive peak per subject (point) for controls and PD patients in response to 3 different feedback types. Stars indicate significance of post-hoc Tukey statistical comparisons (∼ p < 0.1, * p < 0.05, ** p < 0.01, *** p < 0.001). **C**. Scalp topography for each feedback type, centered on the time of the main negative peak (FRN), and for differences with COR FRN (bottom row). On the right, boxplots show the difference between FRN values measured at CO1 and COR and between INC and COR, by groups of subjects and displayed for the midline electrodes.

### Feedback-related oscillations and power spectrum analyses

Neurophysiological markers of feedback processing have been observed consistently at the level of theta and beta oscillations for frontal midline electrodes (Cohen et al., 2007; Marco-Pallares et al., 2008; Marco-Pallarés et al., 2009; Doñamayor et al., 2011; van de Vijver et al., 2011; HajiHosseini et al., 2012; HajiHosseini and Holroyd, 2015). Because such markers are potentially influenced by the systems altered in Parkinson’s disease we also looked for modulations of feedback-related oscillations depending on feedback types and subject groups. Feedback-related activity was thus also studied in the frequency domain.

Time-frequency diagrams revealed differential increases of theta and beta oscillations at the time of feedback for all types of feedback. **Figure 3** shows the significant post-feedback changes in power from baseline for each group of subjects and each feedback type (**Fig. 3A, top**), as well as subject group difference (**Fig. 3A, bottom**). It also shows an analysis focused on specific time windows for theta and beta bands (see methods. **Fig. 3B**). The window analysis showed that theta activity was sensitive to feedback type only (Main effect feedback_type x group: group, F(2,60) = 0.22, p = 0.80, ns; feedback_type, F(2,60) = 11, p = 0.001). Also, for both groups of subjects, feedback-related theta in the first correct trial was greater than in correct trials during repetitions as for the FRN (Post-hoc Tukey contrasts: COR_Control_ – CO1_Control_: Std = -336.3, Error = 94,5, z value = -3.6, p = 0.005; COR_PD_ – CO1_PD_: Std = -253.9, Error = 88.7, z value = -2.9, p = 0.046). Theta for INC differed only marginally from theta in COR trials (F(248.5,94.5) = 2.63, p = 0.086). The refined time-frequency analyses confirmed the absence of subject group effect on post-feedback Theta power (**Fig. 3A, bottom**).

**Figure 3.**
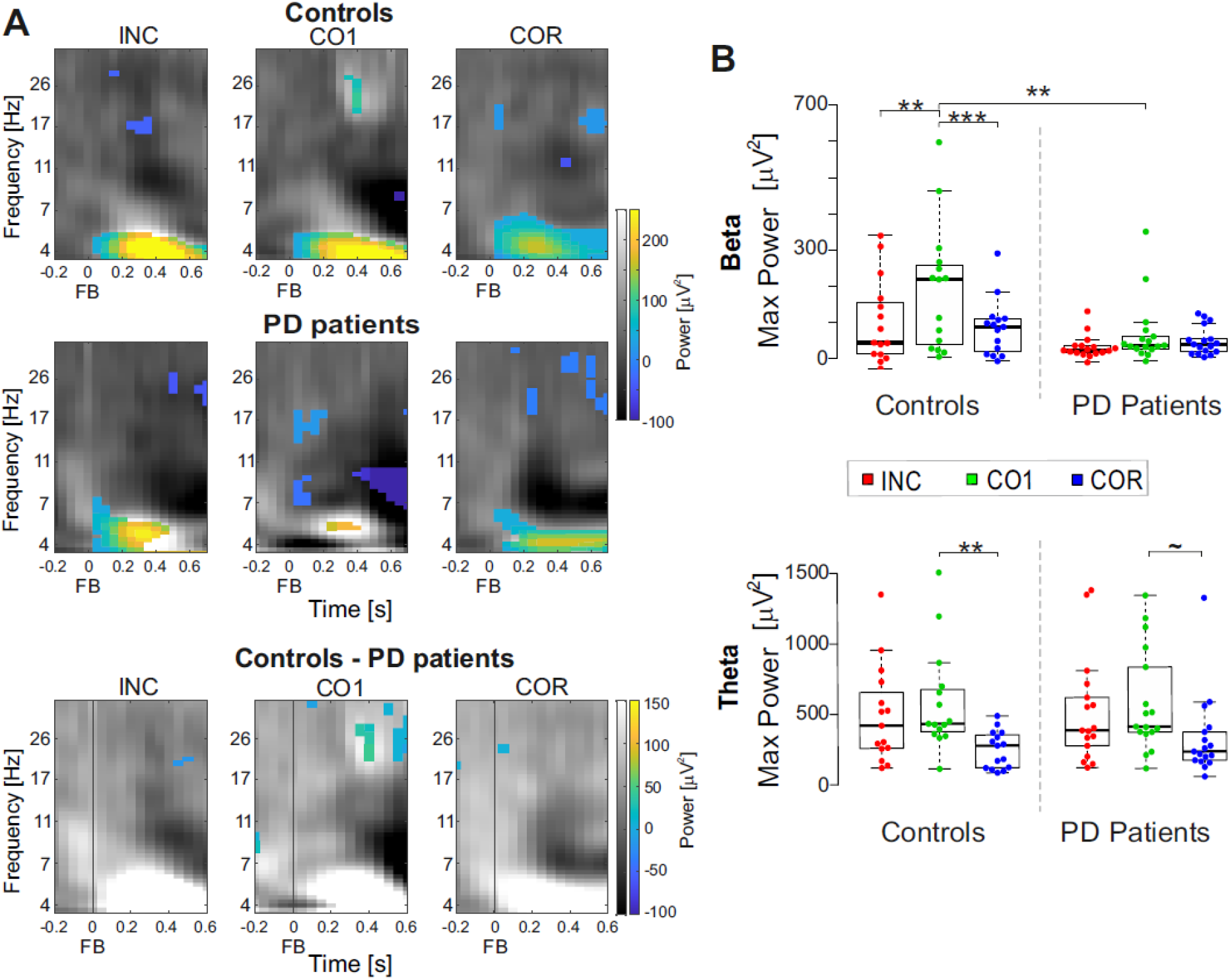
A. Time frequency plots for each feedback type for patients and controls computed for signal recorded at Fz. Top rows. In gray scale the normalized time-frequency plot between minimum (black) and maximum (white), while on the color scale, the power values significantly different from the baseline (−300 to -100 ms prior to feedback), obtained through 1-way ANOVA with FDR correction (p<0.05). Note how theta activity in PD patients loses significant difference evoked by the appearance of feedback compared to baseline, especially in response to the first positive feedback, CO1. **Bottom row**. Time-frequency difference graph of controls minus PD patients for the 3 types of feedback. In gray scale the normalized difference between minimum (black) and maximum (white), while in colors the significant difference between Controls and EP Patients using 1-way ANOVA with FDR correction. Only the beta activity around of 400ms reveals an effect given by the condition (control or PD patient). **B**. Peak-Amplitude distributions per feedback type and group (control, patient) for beta (15 to 30Hz across 300 to 600 ms post feedback) and theta (3 to 7 Hz, across 200 to 600 ms post feedback) bands. Stars indicate significance of post-hoc Tukey statistical comparisons (∼ p < 0.1, * p < 0.05, ** p < 0.01, *** p < 0.001).

In contrast to theta, feedback-related beta power seemed mostly present for control subjects. In the beta range (20 to 30 Hz between 300 to 600ms) the maximum power was significantly altered by feedback type (Main effect feedback type, F(2,60) = 7.46, p = 0.001), and by subject group (Main effect Group, F(1,30) = 7.48, p = 0.01) (**Fig. 3B**). The interaction feedback-type x subject group was marginal (Beta: feedback type x Group, F(2,60) = 2.87, p = 0.064). Further post-hoc analyses showed that increased beta power was indeed observed only in response to CO1 feedback in control subjects, and differed significantly between INC and COR feedback, and compared to CO1 in PD patients (Post-hoc, Tukey Contrast: COR_Control_ – CO1_Control_: Std = -106.17, Error = 26.56, z value = -4, p < 0.001; INC_Control_ – CO1_Control_: Std = -89.73, error = 26.56, z value = -3.4, p = 0.009; CO1_PD_ – CO1_Control_: Std = -122, error = 34.67, z value = -3.5, p = 0.005 ; **Fig. 3B**). Note that since feedback-related changes in power are estimated with reference to pre-feedback power, the results in PD patients (and loss of variance as shown on figure 3B) revealed a complete lack of change in beta power. Contrasting time-frequency decompositions showed between-group effect in the beta range predominantly for CO1 feedback (**Fig. 3A, bottom**). In conclusion, a main effect of group (health condition) was observed for a beta power increase at the time of positive feedback, in particular the 1^st^ positive feedback of problems.

### Spectral changes

Because of the lack of post-feedback beta power increase in patients, we assessed whether more global changes in oscillatory activity could be observed. To do so, instead of focusing on event triggered EEG we analyzed the frequency composition of the 13 first minutes of signal recorded in the session of each subject (including task performance). We focused on the 1 to 60Hz range, and looked in particular within the beta band separating 12-20Hz and 20-30Hz sub-bands. The PSD for one subject stood as an outlier in the analyses and was removed from the group. The average power spectrum clearly revealed the drop in Beta for patients compared to controls, which was not present for theta or alpha bands (One way ANOVAs, Theta (4-8Hz): F(1, 30) = 0.11, p-value=0.74; Alpha (8-12Hz): F(1, 30) = 0.12, p-value= 0.73; Low Beta (12 - 20Hz): F(1, 30) = 4.87, p-value=0.035; High Beta (20-30Hz): F(1, 30) = 7.820, p-value= 0.009; **Fig. 4A**). Neither low nor high beta powers correlated with the individual clinical test scores (Pearson correlations, for PD patients: Low Beta vs DDEL r=0.11, p=0.68; PD: High Beta vs DDEL r=0.06, p=0.84).

**Figure 4.**
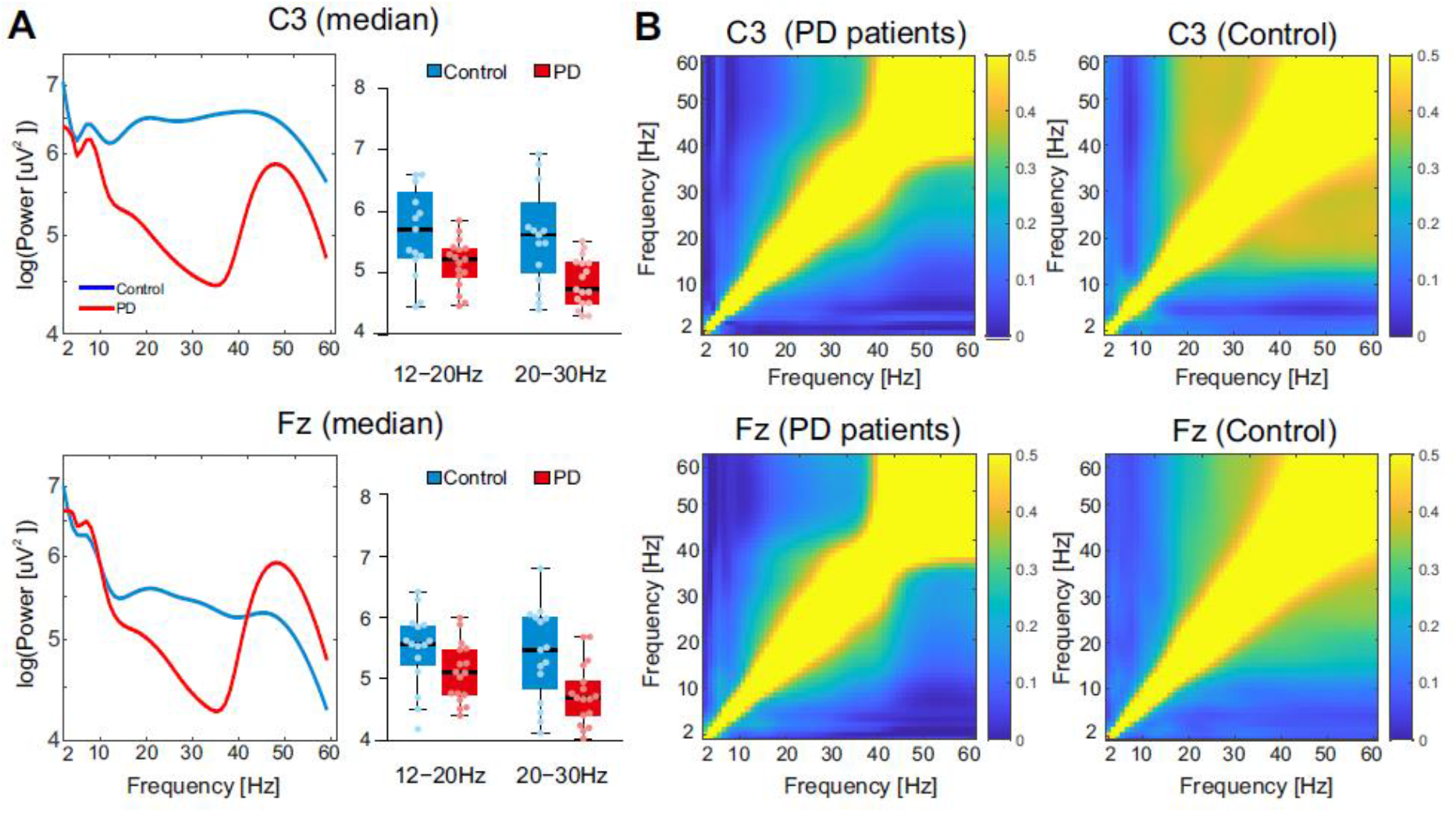
Power spectrum over Fz and C3 in the first minutes of recordings. **A**. On the right, power spectral density of continuous EEG over a period of 13 min for control (blue) and PD patient (red) for signal at electrode C3 (top) and Fz (bottom). On the left, measures for 2 Beta sub-bands for both groups. **B**. Correlation matrices of power spectra of continuous EEG for control (left) and PD patients (right).

As altered cross-frequency correlations have been observed previously in different pathologies (Llinás et al., 1999), suggesting abnormal organization of brain dynamics induced or correlating with pathological conditions, we performed cross-frequency correlations maps specifically for signal captured by electrodes C3 and FZ. The analysis of the continuous EEG over the period of 13 min revealed differences of the cross-frequency correlation pattern in PD patients compared to controls. PD patients showed a lower level of cross-correlation in particular between beta and gamma bands (**Fig. 4B**). The drop in cross correlation is particularly visible on the C3 electrodes, C3 putatively capturing signals from above motor cortex on the contralateral side of the hand used by subjects (Right hand).

## Discussion

We assessed the neuropsychological status, behavioral performance, and EEG markers of performance monitoring functions using a trial-and-error task in recently diagnosed PD patients, compared to healthy controls.

In our study, PD patients presented significantly higher levels of apathy, but not depression, and more cognitive deficits than the control group as evaluated by standard clinical tests. This pattern of neuropsychological impairments in PD patients at early stages of the disease is in accordance with recent studies that encourage a global approach in the research and treatment of Parkinson’s disease, which include the classical motor symptoms but also the non-motor symptoms impacting the cognitive and emotional domains (Foltynie et al., 2004; Aarsland et al., 2009; Kaji and Hirata, 2011; Olanow and Obeso, 2012). At present medical management of PD patients is principally focused on dopaminergic substitution, and neuropsychiatric impairments are frequently underestimated.

The global performance of PD patients in the experimental trial-and-error task was qualitatively similar to that of controls. Patients could understand the task and finished a majority of problems during a session. They nevertheless failed to terminate problems in more cases than controls, and committed more perseverative errors during the repetition period than controls. Such results might be connected to those suggesting executive types of dysfunction in PD especially when subjects are engaged in cognitive tasks requiring active adaptation of decisions and problem solving as measured for instance using the Raven matrices (Brück et al., 2004; Nagano-Saito et al., 2005). RTs were similar in PD patients and controls with higher RTs in search than in the repetition, a pattern that is typically seen in such tasks and that suggest the underlying structure of cognitive progression during problems was not altered in the PD group.

### Feedback-related potentials and oscillations in PD patients

Studying the FRN can provide important clues about the nature of the neural mechanisms contributing to error processing, decision making, and reinforcement learning, both in healthy subjects and patients with alterations of various relevant brain systems as e.g. the dopaminergic system in Parkinson’s disease. One of the aims of this study was to determine if the FRN, modulated by the valence of performance feedback would be impacted in a selective manner by Parkinson’s disease. During the trial and error learning protocol the FRN in control subjects reflected the evaluation of both negative and positive feedbacks when those are relevant for adaptation (i.e. in incorrect trials and for the first positive feedback). An FRN signal at Fz and FCz was elicited by negative feedback during search and to the first reward in a problem. The FRN was reduced for positive feedback during repetition. Our objective was to test whether the early effects of neurodegenerative disease would reveal changes in FRP and possibly for different feedback types (positive versus negative, informative (INC, CO1) versus non-informative (COR). Feedback type effects were not changed in PD patients compared to controls. It somewhat contrasts with a recent study showing that a fronto-central positivity to positive Feedback was diminished in early diagnosed PD patients compared to controls in a probabilistic two alternative forced choice task, but the FRN for negative outcomes (no reward) did not vary between subject groups (Brown et al., 2020). In fact, a differential effect of probability of reward on positive and negative feedback-related potential has been observed in healthy subjects (Cohen et al., 2007).

In the current study we hypothesized that the FRN signal could be altered for the PD patient group after informative feedbacks in particular and potentially differently for positive and negative feedback, which would have been in line with some studies evaluating the ERN in Parkinson’s patients (Willemssen et al., 2009b). The same effect was found for post-negative feedback Theta power increase which also reflects feedback processing (Cavanagh et al 2010 Neuroimage) and has been found altered in Parkinson’s disease patients (Singh et al., 2018). We did not find any significant difference as in some previous studies (e.g. Holroyd et al 2002). The limitations of our study that might explain such negative results are detailed below.

### Parkinson’s disease and task-related oscillations

In controls, theta power and beta power increase were elicited by feedback in search, showing in particular higher increases for the first correct feedback (CO1) compared to positive feedback in repetition. In the context of executive functions and adaptive behavior, theta activity (4 - 8 Hz) has been detected on the frontal midline in response to negative feedback during gambling and learning tasks (Cohen et al., 2007; Marco-Pallares et al., 2008; De Pascalis et al., 2010a), with modulations similar to those for the FRN (Chase et al., 2011; Cunillera et al., 2012; Leicht et al., 2013). Other studies also showed changes of low-frequency activity in response to positive feedback (Doñamayor et al., 2011; HajiHosseini et al., 2012). Moreover, activity at higher frequencies including the beta range between 20 to 30Hz, with a similar frontal midline origin to the theta oscillation, have been described in response to positive feedback (Cohen et al., 2007; Marco-Pallares et al., 2008; Marco-Pallarés et al., 2009; Doñamayor et al., 2011; van de Vijver et al., 2011; HajiHosseini et al., 2012; HajiHosseini and Holroyd, 2015). Cunillera et al. 2012, using a modified version of the Wisconsin Card Sorting Task that includes exploration and exploitation phases comparable to our protocol, reported increased beta activity at the first correct feedback of a series and not during the repetition (Cunillera et al., 2012). Other studies have reported beta activity for negative feedback (Cohen et al., 2008; van de Vijver et al., 2011), or, using magnetoencephalography (MEG) after reward delivery (Doñamayor et al., 2011).

In Parkinson’s disease patients, when compared to the control group, mid-frontal theta activity was not different at the time of feedback compared to other feedback types. Previous reports observed a global change but no differential change in error-related ERP and theta power between error and correct responses in Parkinson’s patients compared to controls (Singh et al., 2018). We found however that post-feedback beta power changes were altered in the PD group especially for positive feedback-related activity in search compared to repetition. Further analyses revealed that beta oscillations were altered (reduced) more broadly, inducing a drop in cross-frequency dynamics involving beta and gamma regimes of oscillations. This result suggests that while feedback-related beta is altered it is unlikely that it reflects a deficit touching selective mechanisms (e.g. feedback-related) but maybe more a global alteration of task-relevant beta power. Abnormal beta oscillations in PD patients have been widely documented (Jenkinson and Brown, 2011; Jackson et al., 2019). Studies have observed an abnormal increase in subcortical beta power as well as in scalp electroencephalography recordings, especially in rest condition (Babiloni et al., 2017; Jackson et al., 2019). In fact, variations of beta power in PD patients are contextually dependant on dopaminergic medication. Studies showed that in the off medication state, EEG inter-electrode coherence in beta frequency increased in PD patients during rest, and the onset of dopaminergic medication reduced it (George et al., 2013). However, during a cognitive task in On-medication state, frontal beta power increased compared to Off-medication, showing that medication reduces abnormal resting beta activity but increases task-relevant beta activity (George et al., 2013). This is in line with reports of reduced anticipatory beta oscillations in rhythmic auditory task, that were recovered with levodopa or STN-DBS while STN-DBS reduced the resting increase in beta oscillation power (Gulberti et al., 2015). In our study, it is likely that the reduced beta power and cross frequency synchrony in the first 13 minutes of task performance was due to such task-related alteration of beta oscillations.

Finally, in a monkey model of Parkinson’s disease, frontal beta power changes have been tracked and partially reflected reduced engagement in the trial and error task induced by a progressive dopaminergic system lesion (Wilson et al., 2016). In the present study, PD patients showed on average significantly worse scores on the apathy scale, but those scores did not correlate with the beta power measured during the first minutes of EEG recordings. Further studies focused on the motivational aspects of the disease and neural markers are required.

## Limitations

This study described feedback-related EEG signals in Parkinson’s patients and healthy controls in a deterministic trial and error learning task allowing analyses over several feedback types. Whereas the study allowed for between-groups comparisons the small sample size in each group must be noted as a limitation for this study. Yet several relevant features of feedback related potentials as well as of feedback-related oscillation power have been observed for both groups and in accordance with past studies. We do not find significant differences in FRN amplitude or theta power between PD patients and controls, but again sample size might have been a limitation. Several studies have also compared FRN and oscillations in PD patients ON versus OFF medication seeking a particular effect of dopaminergic transmission on performance monitoring signals. Unfortunately, the testing conditions for patients at the time and place of the study prevented us to perform such within-subject comparison. Previous studies have observed no effect of medication (OFF vs ON) on error- or feedback-related markers (Singh et al. 2018, Brown et al. 2020, Stemmer et al. 2007) or a decrease of response-related signals under medication compared to control, with no effect of valence (Seer et al., 2017).

## Conclusions

This study shows that some markers of performance monitoring, specifically Beta oscillations, are selectively altered in the early stages of Parkinson’s disease, yet those markers are accompanied by global changes in beta oscillation. Whereas such effects on neurophysiological markers could be relevant for the characterization of the early stages of Parkinson’s it must be noted that these do not correlate with clear behavioral markers or with scores established by clinical tests.

## Acknowledgements

+ This paper is dedicated to the memory of our colleague and friend René Quilodran who recently passed away.

